# Electrostatic features for the Receptor binding domain of SARS-COV-2 wildtype and its variants. Compass to the severity of the future variants with the charge-rule

**DOI:** 10.1101/2022.06.16.496458

**Authors:** Fernando L. Barroso da Silva, Carolina Corrêa Giron, Aatto Laaksonen

**Affiliations:** Universidade de São Paulo, Departamento de Ciências Biomoleculares, Faculdade de Ciências Farmacêuticas de Ribeirão Preto, Av. café, s/no – campus da USP, BR-14040-903 Ribeirão Preto SP, Brazil; Department of Chemical and Biomolecular Engineering, North Carolina State University, Raleigh, NC 27695, United States; Universidade Federal do Triângulo Mineiro, Hospital de Clínicas, Av. Getúlio Guaritá, 38025-440 Uberaba, MG, Brazil; Department of Materials and Environmental Chemistry, Arrhenius Laboratory, Stockholm University, SE-106 91 Stockholm, Sweden; State Key Laboratory of Materials-Oriented and Chemical Engineering, Nanjing Tech University, Nanjing, 210009, PR China; Centre of Advanced Research in Bionanoconjugates and Biopolymers, Petru Poni Institute of Macromolecular Chemistry, Aleea Grigore Ghica-Voda, 41A, 700487 Iasi, Romania; Department of Engineering Sciences and Mathematics, Division of Energy Science, Luleå University of Technology, SE-97187 Luleå, Sweden; University of Cagliari, Department of Chemical and Geological Sciences, Campus Monserrato, SS 554 bivio per Sestu, 09042 Monserrato, Italy

**Keywords:** electrostatic interactions, pH effects, molecular simulation, complexation, pKa, virus, electrostatic coupling

## Abstract

Electrostatic intermolecular interactions are important in many aspects of biology. We have studied the main electrostatic features involved in the interaction of the receptor-binding domain (RBD) of the SARS-CoV-2 spike protein with the human receptor Angiotensin-converting enzyme 2 (ACE2). As the principal computational tool, we have used the FORTE approach, capable to model proton fluctuations and computing free energies for a very large number of protein-protein systems under different physical-chemical conditions, here focusing on the RBD-ACE2 interactions. Both the wild-type and all critical variants are included in this study. From our large ensemble of extensive simulations, we obtain, as a function of pH, the binding affinities, charges of the proteins, their charge regulation capacities, and their dipole moments. In addition, we have calculated the pKas for all ionizable residues and mapped the electrostatic coupling between them. We are able to present a simple predictor for the RBD-ACE2 binding based on the data obtained for Alpha, Beta, Gamma, Delta, and Omicron variants, as a linear correlation between the total charge of the RBD and the corresponding binding affinity. This “RBD charge rule” should work as a quick test of the degree of severity of the coming SARS-CoV-2 variants in the future.

Categories and Subject Descriptors:

## 1. Introduction

Viruses are generally exciting systems to investigate electrostatic interactions. In his classic paper on electrostatic effects in proteins, Perutz identified how these interactions played an important role in triggering the assembly of proteins from the tobacco mosaic virus.^1^ This example is just one of the countless processes involving viruses that use electrostatic interactions. There are many more that could be mentioned (see ref. ^2^ for a review). Host-pathogen interfaces contain more titratable amino acids than other protein-protein interfaces (especially TYR).^3,4^ Epitopes have already been discussed within a broader perspective of “electrostatic epitopes” (EE).^5^ This is due to the long-range nature of electrostatic interactions and the possibility of high electrostatic coupling between ionizable residues spread in the protein’s interior and not only at the protein surfaces.^5^ Since the early days of computational protein electrostatics with the pioneer contributions from Åqvist, Sharp, and others,^6,7^ different authors have also shown how important are electrostatic interactions for optimal antibodies.^8–11^ Studies have demonstrated the significance of electrostatic interactions in the RNA-binding specificity and transactivation activity for human immunodeficiency virus (HIV) and simian immunodeficiency virus.^12^ It was also been shown that efficient drugs binding to HIV require specific electrostatic features.^13^ Moreover, pH is well known to trigger several key biological events for viruses. For instance, a drop in pH is often used to allow the host penetration.^14–16^ The influenza virus, for example, needs the low pH inside the endosomal compartment to fuse its viral membrane with the endosomal membrane to release the viral genome into the cytoplasm.^17^ Such a mechanism can also induce changes in the extracellular pH.^18–20^ Flaviviruses such as the Zika virus rely on pH to control the self-association of its NS1 protein which is intimately related to virulence.^21^ Another common example is the homodimerization of the envelope protein of flaviviruses.^22^ Therefore, it is not a surprise that SARS-CoV-2, the virus responsible for the COVID19 pandemic, would have interesting and particular electrostatic features.

A large number of papers have shown that electrostatic interactions are indeed very important for SARS-CoV-2 (e.g. refs. ^23–36^). Most of the studies have focused on the interactions involving the receptor-binding domain (RBD) of the SARS-CoV-2 spike homotrimer glycoprotein due to its biological central relevance. In particular, the RBD interaction with the human cell receptor Angiotensin-converting enzyme 2 (ACE2) and possible binders have been exhaustively explored in the literature both for the wild-type (wt/Wuhan) virus and for its variants. ACE2 is the main cell entry mechanism of the virus.^37^ Without the RBD-ACE2 binding, no infection would occur. Also, this is consequentially the biological interface (see Figure 1) more often investigated for vaccine developments.^38,39^

**Figure 1.**
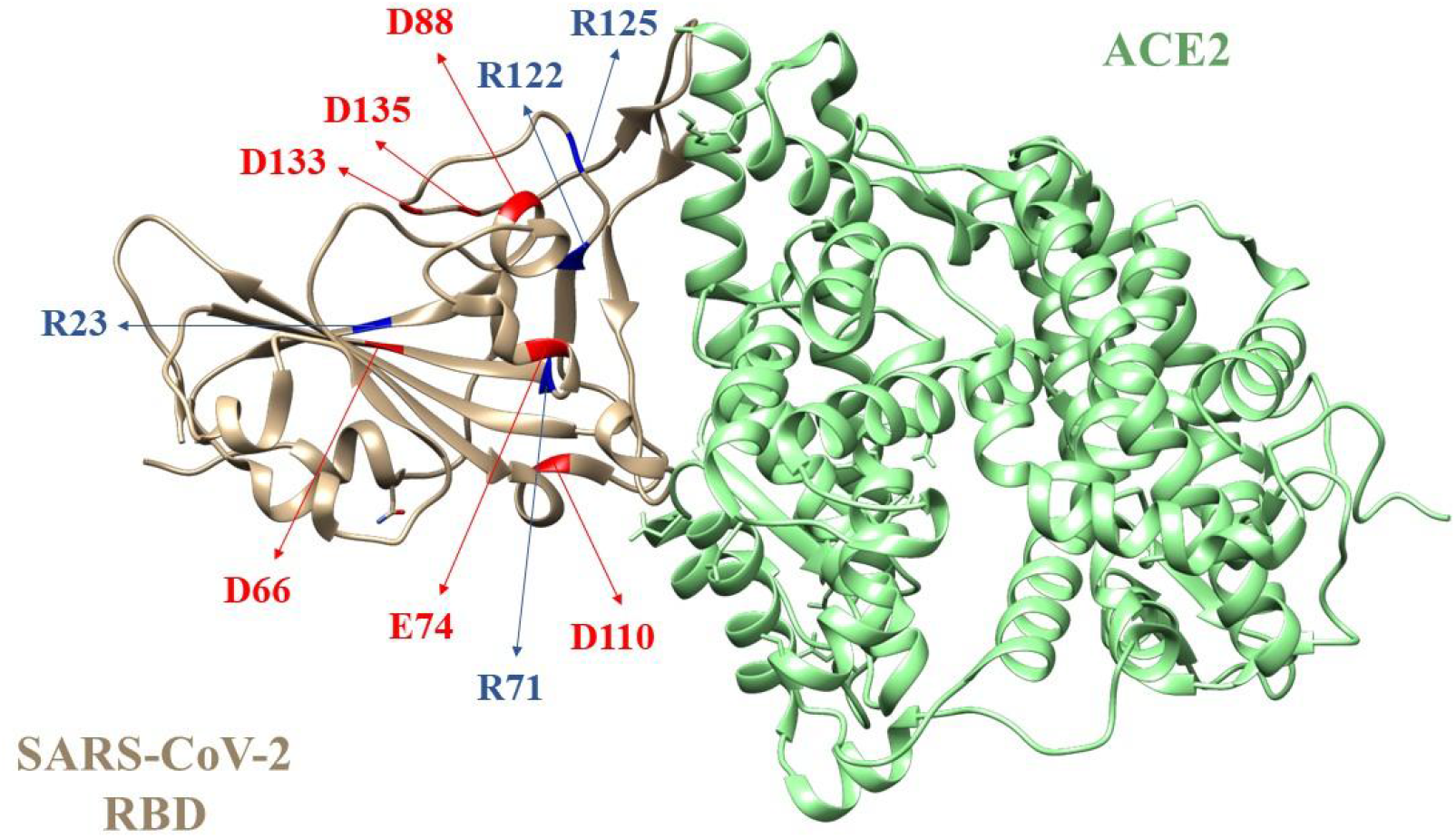
The molecular structure of the SARS-CoV-2 S RBD wt complex with ACE2. These macromolecules are shown in wheat and green, respectively. Some key amino acids of the RBD wt with a high electrostatic coupling as calculated in this work are indicated with red (acid residues) and blue (basic residues) labels. The RBD is also tyrosine-rich.^40^ See text for more details.

From the theoretical perspective (in chronological order), the RBD-ACE2 and RBD-binders biological interfaces for the SARS-CoV-2 wt were investigated at the beginning of the COVID19 pandemic by Giron and colleagues^9^ and followed by several other works (e.g. refs. ^24,41–49^). In the study carried out by Giron and co-authors, it was demonstrated using a constant-pH coarse-grained model and Monte Carlo (MC) simulations that the RBD of SARS-CoV-2 wt could bind to ACE2 with similar binding affinities to SARS-CoV-1. Yet, the ACE2 interaction with SARS-CoV-2 RBD wt involves more titratable groups than its ancestral as later it was also seen in experimental data.^50^ The involved EEs were mapped via the PROCEEDpKa approach^51^ confirming this viral entry mechanism and highlighting the importance of electrostatic interactions for this process.^9^ Four initial known monoclonal antibodies (mAb) for SARS-CoV-1 (80R, CR3022, m396, and F26G29) were tested and a new optimized binder based on CR3022 was suggested based on the Coulombic interactions.^9^ In both the RBD-ACE2 and RBD-mAbs studied systems, electrostatic interactions were clearly important. This has been confirmed for both the SARS-CoV-2 RBD wt [RBD_(wt)_] and the subsequent variants by many other theoretical studies that employed different molecular models, a diversity of simulation techniques, and also input protein structures (different atomistic coordinates from the RCSB Protein Data Bank and/or from homology modeling). For instance, a couple of days after this pioneering work, Spinello and collaborators dissected the key molecular interactions for the RBD_(wt)_-ACE2 complexation using microsecond-long all-atom explicitly solvated molecular dynamics (MD) simulations (Parm99SB-ILDN AMBER force field [FF]). Again, the importance of electrostatic interactions for this process was observed.^52^ Almost at the same time, another group integrating a 300 ps MD (AMBER-f99SB FF) simulations and a Poisson-Boltzmann (PB) solver combined with an MC scheme reported the electrostatic potentials for SARS-CoV-1 and 2 RBD_(wt)_ proteins (more positively charged), and ACE2 (more negatively charged), mapped the key salt bridges for the complexation and described the enhanced electrostatic interactions observed for SARS-CoV-2 RBD_(wt)_-ACE2 interaction.^53^ Steered all-atom MD (AMBER-f99SB-ILDN FF) and coarse-grained (a Go-like model) simulations carried out by Nguyen and co-authors discussed the lack of consensus in the experimentally measured binding affinities, reproducing the experimental trends reported by Walls et al.^54^ and provided more data to increase the consensus that the RBD_(wt)_-ACE2 complexation was driven by electrostatic interactions.^48^ Bai and Warshel also indicated the importance of electrostatic interactions^43^ but supported the idea that residues located outside the RBD_(wt)_ were critical for the complexation. A multi-scale computational approach combining PB calculations and 50 ns of MD simulations demonstrated the attractive electrostatic forces between the RBD_(wt)_ and ACE2 at different distances.^55^ They have shown as well that SARS-CoV-2 RBD_(wt)_ has more high-occupancy hydrogen bonds at the RBD_(wt)_-ACE2 interface area than SARS-CoV-1 RBD which agrees with above mentioned theoretical and experimental data. Protein-protein docking studies pointed out in the same direction.^56^ There is undoubtedly well-built support for the importance of electrostatic interactions for this process which is constantly confirmed for the latest variants, ACE2 polymorphisms, and new studied Abs. ^23,32,36,57–62^ Nevertheless, it is worth mentioning that Jawad and co-authors more recently added that the van der Waals interactions can also be critical to stabilizing the RBD-ACE2 complex even if the Coulombic interactions are the main driving force for the recognition process between the RBD and ACE2.^59^ Although the main aspects are consistently reproduced, quantitative results for the predicted binding affinities can vary a lot as was critically discussed by Taka and colleagues.^63^

Despite an incredibly high and continuously growing number of works on this research topic that is evolving so fast, there are still open questions and preliminary results that need to be revisited particularly due to the lack of a systematic prediction of all of them on the same basis and by the same theoretical tools. Most of the mutational studies focused on hydrophobic residues (not on titratable groups) and/or specifically on the ones located in the RBD-ACE2 binding interface.^64,65^ Although it is clear that pH can have a crucial role in the interaction between the RBD and its biological partners, the majority of the theoretical studies were performed with constant-protonation states simulations (i.e. the charges of the amino acids are defined at the input phase for a given pH condition and remain constant during all the simulation run) instead of CpH ones (i.e. the protonation states are constantly updated during the simulation run for a given input pH condition. Here, pH is an input parameter as is the temperature).^66^ This implies that possible contributions from mesoscopic electrostatic mechanisms as the charge regulation phenomena due to the proton fluctuations^67–69^ were not considered and/or quantified in many previous works. Similarly, calculations were performed almost exclusively at pH 7. This can be of limited use for some recent pharmaceutical applications. The second generation of vaccines and treatments might need to explore other alternative routes where pH can play again important role. For instance, vaccines via nasal spray that could activate the so-called mucosal immune system^70,71^ (see also clinical trial NCT04954287) would need to deal with larger pH windows and not only data at pH 7. This is also the case for intranasally-delivered antibodies.^72^ However, the experimental data for the nasal mucosal pH seems to be quite scattered. Values have been reported within the pH range of 5.3-7.0.^73^ Patients with rhinitis can have higher values (7.2-8.2).^73^ Gender and ethnic differences can also affect the nasal pH.^74^ A description of the pH effects of the RBD and its interactions would provide useful physical-chemical insights for such developments.

Motivated by such issues, we performed an extensive set of simulations to investigate the details of the electrostatic features of the RBD for the several variants, their main physical-chemical properties (net protein charge, charge regulation capacity, dipole moment, and the *pKas* for all ionizable residues), binding properties, and how these physical quantities are connected at several pH regimes. The contributions from Coulombic, charge regulation, and dipole interactions were dissected. The electrostatic coupling between titratable groups was mapped. Comparing different viral variants employing the same theoretical tools and physical-chemical conditions, the present study reveals how the virus evolution is exploring these electrostatic features to improve the RBD-ACE2 binding affinity which impacts the virulence, transmission, and infectivity. Together with more insights on the importance of electrostatic interactions for viruses, the present work also contributes to understanding the trajectory of severe acute respiratory syndrome coronavirus 2, its evolutive processes, and possible correlations with the virus transmissibility and infectivity. Based on the present analysis, we proposed as well a simple predictor for the ACE2 binding affinity using titration properties that can be useful to estimate the degree of severity for future mutations.

## 2. Model and methodology

This study requires an extensive set of simulations with a large number of runs to investigate the details of the principal electrostatic features of the RBD for the wt, several variants (Alpha,^75^ Beta,^76^ Gamma,^77^ Delta,^78^ and Omicron^79^) and sub-variants that recently appeared as a threat (BA.2, BA.3, BA.4, BA.5, and BA.2.12.1) under different physical-chemical conditions. In particular, the estimation of the free energy of interactions for protein-protein interactions by computer simulations is well known to be highly computational demanding. This situation naturally justifies approximations invoking molecular models with a reduced degree of freedom and consequentially a smooth energy landscape. At the same time, it is expected that such coarse-grained (CG) simulations of biomolecules should accurately describe the electrostatic interactions and pH effects. Among possible available tools,^80–82^ the FORTE (**F**ast c**O**arse-grained p**R**otein-pro**T**ein mod**E**l)^10,21,23^ approach was a suitable choice to fulfill all these previous issues and also due to its record of successful applications in other biophysical systems^67,83–85^ (including viruses^9,10,21,23,32,51,86^) where electrostatic interactions are quite important. This was the main computational tool used in the present work. It is a constant-pH (CpH) CG model that incorporates proton fluctuations and combines it with random translations, and rotations of rigid protein models solved by Metropolis Monte Carlo (MC)^87^ to compute the free energy of interactions [*w(r)*] for protein-protein systems. Despite its intrinsic approximations, FORTE can reproduce the main features of more sophisticated methods with lower CPU costs.^10^ Conversely, each individual simulation is not necessarily cheaper and still demands a lot of high-throughput computing. The number of systems and conditions investigated easily resulted in a large ensemble of simulations (here, more than 58k runs were performed combining resources from different computer centers). They also often require a large number of MC steps due to the energetic barriers of the systems and the need to record the frequencies for the two proteins at different separation distances (*r*) with a reasonable number of observations. A cylindrical simulation box is used to facilitate the histogram sampling employed to estimate *w(r)*. This is different from traditional docking procedures that focus only on short-range interfaces and often do not allow changes in the protonation states during the runs. Here, an ensemble of equilibrium configurations with the simulated systems at different protein-protein separation distances, orientations, and protonation states are obtained and analyzed to estimate *w(r)* for each studied system and physical-chemical condition.

Such an approach is conveniently applied in a large-scale scenario context where a very large number of simulation runs are needed. The FPTS (**F**ast **P**roton **T**itration **S**cheme)^88,89^ is the engine responsible to control the acid-basic equilibrium of all titratable groups at each studied pH condition. This titration scheme for proteins and nucleic acids combines a good accuracy, a reasonable physical description of the system (with the possibility of describing salt effects and external electric fields), and a lower CPU cost in comparison to Poisson-Boltzmann solvers and other CpH methods.^66,88–90^ A drawback of this approach is that hydrophobic “interactions” are only partially taken into account.^83,85^ Mutations involving only non-ionizable groups (e.g. LEU→ALA) might also be not so well described as the titratable ones. Conversely, the model is ideal to explore electrostatic interactions as is the aim of this study.

### Molecular systems and their modeling

The present work focus exclusively on the second step of the two-step “expose–dock-like” mechanism for the complexation RBD-ACE2.^9,23,32^ Therefore, it is assumed here that the RBD was available to interact with the ACE2 with no steric clashes with other parts of the spike (S) homotrimer. This would correspond to the S protein being all the time at the “up” conformational state.^9,23,32^ The simulated model does not incorporate the transition between the “up/down” states of the S protein. Only the RBD of the S protein is present in the simulations. All studied proteins (RBD and ACE2) were modeled at the CG level as a set of spherical charged Lennard-Jones (LJ) beads mimicking their amino acids. Charges of the titratable groups (*z*_*i*_) are pH-dependent and can vary during the simulation runs depending on the environment. They are assigned and updated during the runs as a function of the input pH by the FPTS. In doing so, the electrostatic interactions are fully described by this simple and robust model. The charge regulation mechanism that can result in additional mesoscopic attractive electrostatic forces^68,69,91^ is properly incorporated. Non-titratable amino acids are kept neutral all the time. The size of the LJ beads (*R*_*i*_) is taken from ref. ^92^. This possibility to treat each residue type by a different value of *R*_*i*_ is a simple way to partially account for the non-specific contributions from the hydrophobic effect in the model.^10,51,83^ Such different radii help to both preserve the macromolecular hydrophobic moments and to guide the orientation at short separation distances. Other details, justifications, validation, pros, and cons are given elsewhere.^9,51,83,85^ In short, the full configurational energy of the system [*U*({*r*_*k*_})] combining a screened Coulombic term (first term) with an LJ one (second term) is given by

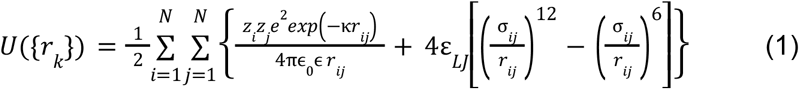

where *{r*_*k*_*}* are amino acid coordinates at a given configuration, *z*_*i*_ and *z*_*j*_ are the valences of the amino acids *i* and *j* as defined by the titration scheme, *e* is the elementary charge (e=1.602×10^−19^C), κ is the modified inverse Debye length,^88,93^ *r*_*ij*_ is the separation distance between beads *i* and *j, ε*_*0*_ is the dielectric constant of the vacuum (ε_*0*_ =8.854×10−12 C^2^/Nm^2^), *ε* is the dielectric constant of the medium assumed to be equal to 78.7 to mimic an aqueous solution at temperature *T* equals to 298K, *ε*_*LJ*_ is a parameter that regulates the attractive forces in the system,^51,83^ *σ*_*ij*_ is given by Lorentz-Berthelot rule from the radius of beads *i* and *j* (*R*_*i*_ and *R*_*j*_), and *N* is the total number of beads. The universal value of 0.124 kJ/mol was chosen for *ε*_*LJ*_.^83,85^ This should be equivalent to a Hamaker constant of ca. 9 *K*_*B*_T (*K*_*B*_ is the Boltzmann constant) for amino acid pairs.

Following the previous works,^9,23,32^ the coordinates of the ACE2 molecule were obtained from the PDB id 2AJF (chain A, resolution 2.9 Å, pH 7.5). The SARS-CoV-2 RBD_(wt)_ coordinates were taken from ref.^9^ to allow a proper comparison with recent studies. The coordinates are also available at Zenodo.^94^ Structural comparisons between the RBD_(wt)_ and some mutations have shown that the molecular sequence-structure relationship is very powerful to preserve the overall folded structure.^79^ This has been surprisingly observed even for Omicron with 15 amino acid replacements at the RBD - see Table 1. Therefore, mutations were modeled by the simple replacement of residues from the wt protein followed by an energy minimization using “UCSF Chimera 1.14”.^95^ Rotamers were created using the Dunbrack 2010 library. The ones with the highest probabilities were selected for each mutation. Other default parameters were chosen for the minimization procedure. A list of variants and the corresponding replacements are given in Table 1. The ratio of the number of the titratable groups involved in these mutations is also listed in this table. In most cases, either the mutation happens in one ionizable amino acid (e.g. **K**417N) or a non-titratable one is replaced by an ionizable one (e.g. Q493**R**). The majority of mutations are related to titratable amino acids anticipating the role of electrostatic interactions. Even variants with the *R*_*titra*_ < 1 (as seen for Omicron) involve a larger number of titratable residues than the ones with *R*_*titra*_ equal to 1. In Omicron, 9 mutations (G339**D, K**417N, N440**K**, T478**K, E**484A, Q493**R**, Q498**R**, N501**Y**, and **Y**505**H**) out of the 15 involve ionizable amino acids (1 D, 3 K, 1 E, 2 R, 2 Y, and 1 H). On the other hand, the Delta variant has 3 mutations (L452**R**, T478**K**, and **E**484Q), and all of them contain titratable groups.

**Table 1.**
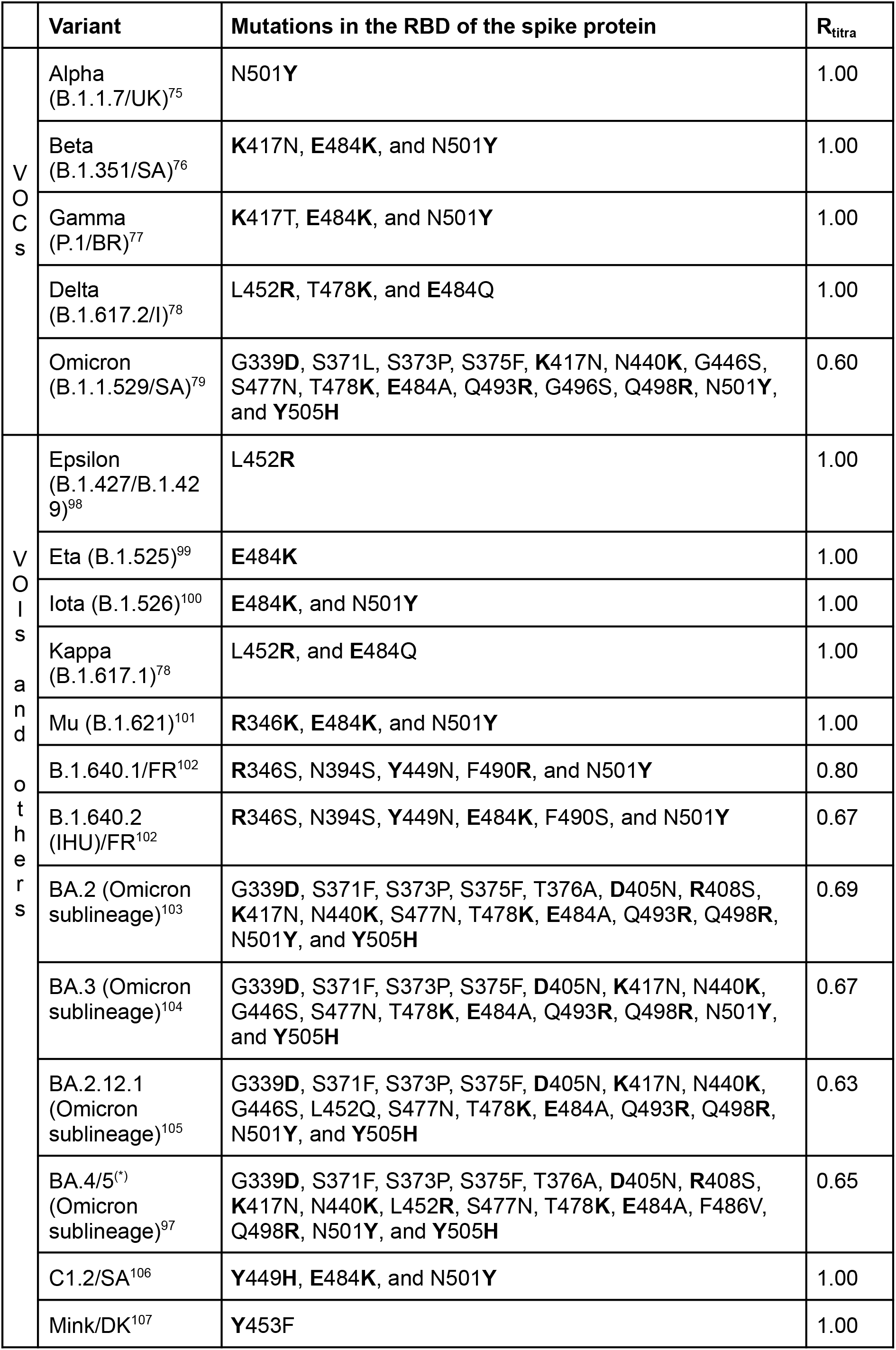
Mutations in the RBD of the spike protein on each studied variant. Titratable residues are highlighted in bold. VOCs and VOIs are, respectively, the “variants of concern” (mutations that are circulating more than others, causing more severe disease, or are more transmissible) and “variants of interest” (mutations that are circulating more than others with a risk to be upgraded to a VOC) as classified by the World Health Organization.^96^ BR, DK, FR, I, SA, and the UK refer, respectively, to Brazil, Denmark, France, India, South Africa, and United Kington, i.e. the countries where these variants were first detected. **R**_**titra**_ is the ratio between the number of mutations involving at least one ionizable amino acid and the total number of mutations. ^(*)^ The variants BA.4 and BA.5 share the same mutations in the protein S. Yet, they have different sets of mutations in the other proteins from the remaining genome.^97^

### Molecular simulation details

The RBD-ACE2 complexation calculations were performed within the pH range of 1 to 14 at physiological ionic strength (150 mM of NaCl) and using a simulation cylindrical cell with a radius of 150Å and height equals 200Å. Averaging was carried out over an additional 3.10^9^ MC steps using the Metropolis algorithm after a proper equilibration. The frequency that the RBD and ACE2 molecules were observed at a separation distance between *r* and *r*+*Δr* was recorded in a histogram during this production phase. The histogram bin size *Δr* was 1Å. After the appropriate normalization, the histogram was converted to a pair radial distribution function [*g(r)*]. The free energy of interaction for the complexation process was calculated from the usual relation: *w(r)=-ln[g(r)]*, where is equal to the reciprocal of *K*_*B*_*T*.^108^ Standard deviations were estimated using three replicates per simulated system. Additional simulations with only one protein in an electrolyte solution were also performed to provide the average charge of each amino acid (⟨*Q*_*i*_⟩ *=* ⟨*z*_*i*_⟩*e*), the average total protein charge (⟨*Q*⟩ *=* ⟨*Z*⟩*e*, where ⟨*Z*⟩ is the average total valence), average total dipole moment (⟨*μ*⟩), and the average charge regulation capacity (⟨*C*⟩) at all pH values. *pKas* were calculated from the values of ⟨*Q*_*i*_⟩ obtained at pH solutions from pH 1 to pH 14 using an interval of 0.1. See the ref. ^88^ for more details. The contribution of individual titratable amino acids in the ACE2 binding was mapped through a computational Alanine scanning where each residue is replaced by ALA and the calculations are repeated with this mutation. ^9^

## 3. Results and discussion

### Physical chemistry properties of the proteins

To gain insights into the complexation of two biomacromolecules and the energetic contributions to the free energy of their interaction, certain key physical-chemical properties (*Z, C*, and *μ*) are very useful. Figure 2 displays these properties as a function of pH for the RBD_(wt)_, the five VOCs, and three recent sub-variants of Omicron. The average net charge numbers are given in Figure 2a. All studied systems have qualitatively similar plots.

**Figure 2:**
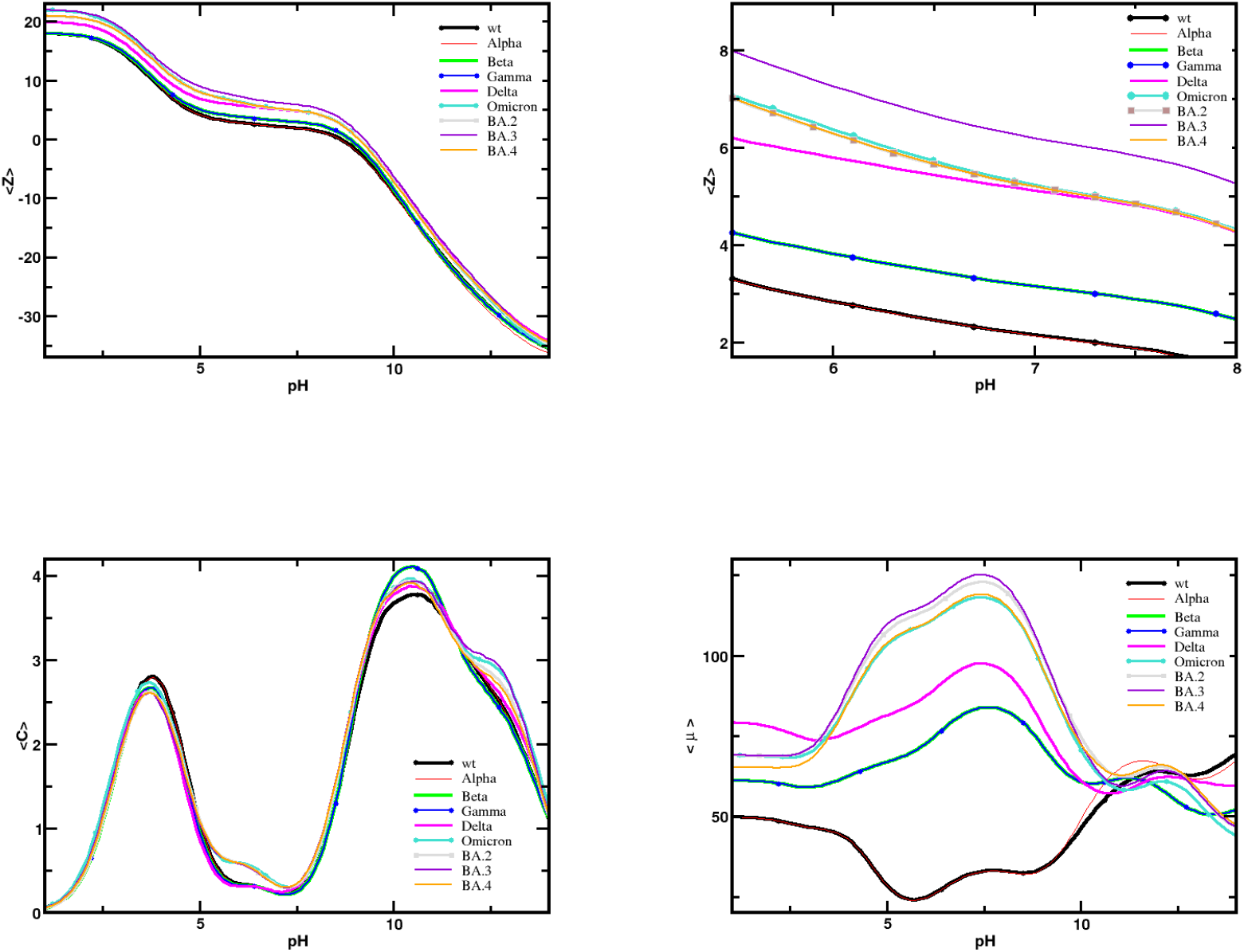
Main physical-chemical properties of the SARS-CoV-2 RBD from the wild-type and different variants. (a) Simulated net protein charge number as a function of pH. (b) The same as (a) with zoom in the pH window 5.5-8.0. Symbols used in the plot were increased to facilitate the comparisons. (c) Simulated charge regulation capacity as a function of pH. (d) Simulate dipole moment numbers as a function of pH. Data from MC simulations at 150 mM of salt concentration. We note that some curves are on top of each other.

Nevertheless, the curves are shifted upwards for all the variants except for Alpha whose titration data is very close to the wt. The data for BA.5 (not shown in Figure 2) is identical to BA.4 due to their sequences (see Table 1). Newer variants (e.g. BA.3) are more positively charged in both the acid and basic pH regimes in comparison with the wt and older variants (e.g. Alpha). The difference decreases upon the increase in the pH. The plots for Beta and Gamma variants are virtually on the top of each other as it is expected. Their difference at the sequence level has involved the same type of physical-chemical replacement (the basic **K**417 is substituted by N or T - both N and T are non-titratable amino acids). Omicron (BA.1), BA.2, and BA.4/BA.5 behave in the same way in terms of their Coulombic contributions. BA.3, a quite recent sub-lineage of Omicron, is the variant that stands out and where larger differences are observed in its titration plots when compared to the others. It is easier to see these comportments in Figure 2b where zoom is given to the pH window between 5.5 and 8.0. At pH 7, ⟨*Z*⟩ is equal to 6.2 for BA.3 while it is 5.2 for Omicron, BA.2, and BA.4/BA.5. BA.2.12.1 (data not shown in this Figure) has a titration plot identical to BA.2. The two variants only differ by the mutation L452Q presented in the BA.2.12.1 and absent in BA.2 (see Table 1) which does not alter the acid-basic equilibrium as captured by our theoretical model. This increase in the net charges has biological implications and also affects how the virus might interact with surfaces (e.g. electret fiber of the masks).^35^ Similar results and analysis at pH 7 are found in the recent paper by Gan and colleagues using the Poisson-Boltzmann approach (except for BA.4 and BA.5 which were not included in their study).^62^

The order of the computed isoelectric points (pI) is wt, Alpha (8.6) < Beta, Gamma (8.9) < Delta, Omicron, BA.2, BA.2.12.1, BA.4/BA.5 (9.1) < BA.3 (9.3). From the computed titration plot for ACE2, its pI is 4.8 at 150mM of NaCl [at pH 7, <*Z*_(ACE2)_>=-23.1(1)]. It becomes evident the attractive Coulombic interactions between the RBD and ACE2 between the pH window of 4.7 and 8.6-9.3 depending on the variant. The net charges for the RBD_wt_ [<*Z*_(RBDwt)_>=+2.2(1)] and ACE2 [<*Z*_(ACE2)_>=-23.1(1)] at pH 7 are comparable with the results from calculations using the ab initio fragment molecular orbital method (<*Z*_(RBDwt)_>=+2 and <*Z*_(ACE2)_>=-24).^36^

The charge regulation capacity for these proteins is given in Figure 2c. Two peeks can be seen: a smaller one at pH 3.6-3.8 with <*C*>∼2.6-2.9 and a higher one at pH 10.4-10.6 with <*C*>∼4.7-4.1. Around physiological conditions (pH 7), the values are smaller than 0.4 which suggests a minor role of the charge regulation mechanism in this pH regime. Yet, it is interesting to see that the more recent variants as the Omicron family started to have higher values of *C* in comparison with the older variants. It could be an indication that the virus evolution is exploring this physical property to increase its binding affinity. A new feature that can have an impact not only on their interactions with ACE2 but with any Ab or other charged molecule. This trend becomes stronger for the dipole interactions despite the sequence identity between these RBD proteins. As it can be seen in Figure 2d, there is a larger increase in the values of *μ* for the newer variants in particular around pH 7. In general, *μ* gradually increased for each new variant that appeared following the time-line they started to circulate. The RBD_(BA.3)_ has a dipole moment number of 124 while the RBD_(wt)_ has only 29 at pH 7 and 150mM of salt, i.e. the *μ*[RBD_(BA.3)_] is surprisingly 4.3 times larger than *μ*[RBD_(wt)_]. It is impressive that virus evolution has been able to tune all these physical properties to increase its attractiveness to ACE2. To our knowledge, it is the first time that such mechanisms involving all these three electrostatic properties have been verified for coronaviruses. Similar behavior was reported for the electrostatic properties of the NS1 protein of the Zika virus from different strains.^109^

### pKa calculations

More detailed information on the source of the different physical-chemical properties above discussed can be found in the individual contributions from each titratable residue. To identify it, we calculated *pKa* values for the RBDs for the wt, the VOCs (Alpha, Beta, Gamma, Delta, and Omicron), four recent variants from the branches descending from an original Omicron ancestor (BA.2, BA.3, and BA.4/BA.5), and the 12th lineage to branch off from BA.2 (BA.2.12.1). As mentioned above, the values for BA.5. are the same ones computed for BA.4 because they share an identical RBD profile.^97^ Table 2 lists their *pKa* values as calculated by the FPTS at 150 mM of salt. As far as we are aware, there is no experimental data nor other theoretical published data for comparison. We rely on the fact that the *pKa*s computed by the FPTS are usually in good agreement with experimental data both for proteins and RNAs.^66,85,88–90^ The ionizable groups that are different by 0.2 *pKa* units from their values in the wt protein sequence are highlighted in bold. The *pKa* values for several groups are relatively stable across all studied variants. This suggests that these particular residues experience similar electrostatic environments despite the differences in their sequences. Nevertheless, note that the number of amino acids marked in bold increases for the newer variants suggesting they have different local electrostatic environments and consequentially distinct interactions. Computed *pKa* for the variants can have shifts from -0.3 to 0.9 pH units from the values obtained for the wt protein. For some RBD proteins, this results in major differences between their computed values and the ideal *pKa* ones. For example, ARG-403 [*pKa*(Alpha)= *pKa*(Beta)=*pKa*(Gamma)=14] experienced a shift of 2.0 pH units in comparison to the ideal *pKa* of 12. Arginines are the amino acids whose *pKa* differs more from their ideal values. The other ionizable groups including the histidine have smaller maximum differences (∼ 1 pH unit). For tyrosines (ideal *pKa*=9.6), there is a more peculiar behavior. It exhibits up and down-shifts with the smaller maximum differences in both directions (0.5 and -0.4, respectively): TYR-501 (an amino acid that appeared mutated in some variants) has *pKa*(BA.2)=*pKa*(BA.4) =*pKa*(BA2.1.21)=9.2 while TYR-365 has *pKa* values for all RBD proteins ∼10.1.

**Table 2.**
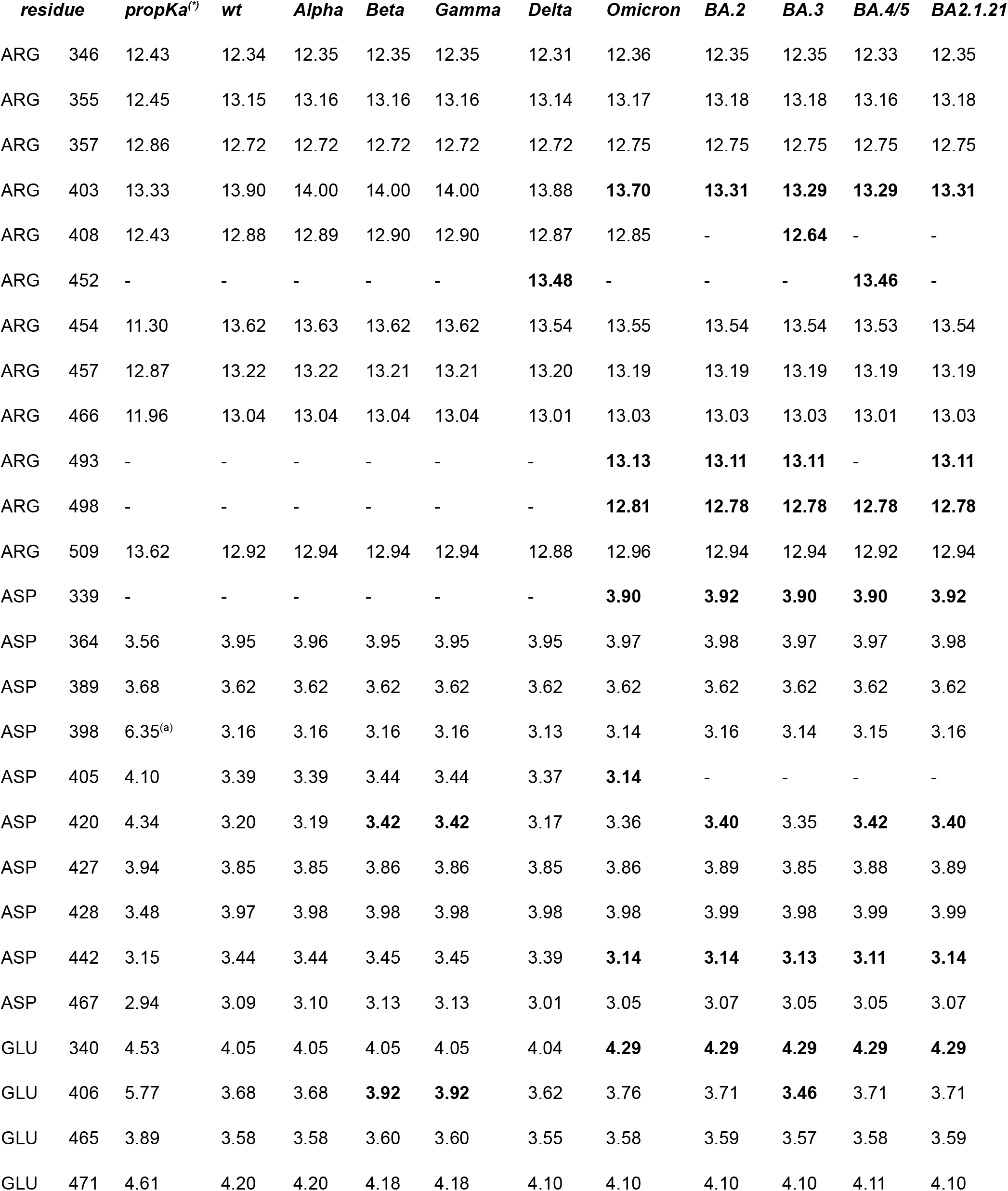
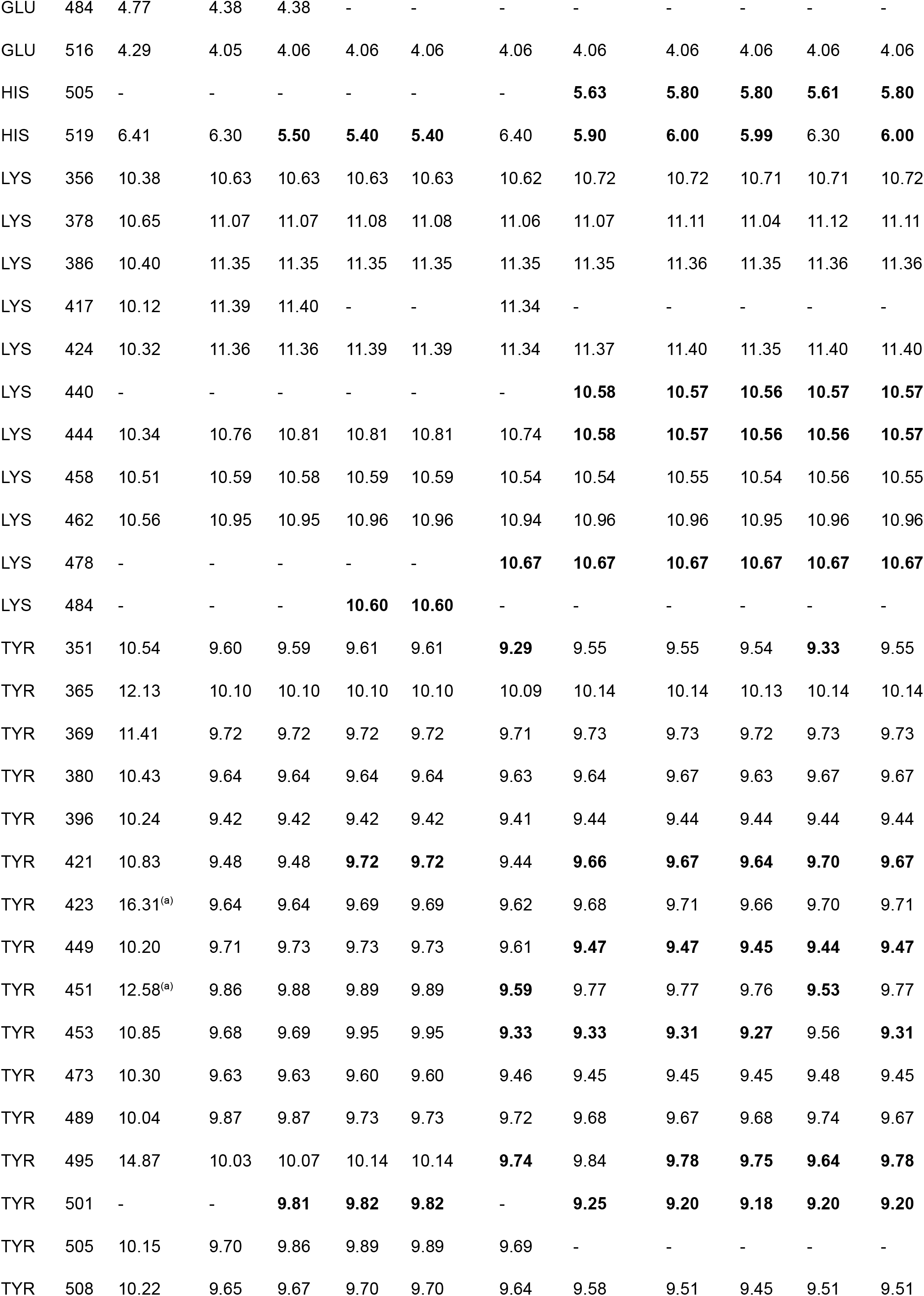
Calculated *pKa* values for the RBD wt and several mutations at 150 mM of salt. Data from the FPTS at 150mM of NaCl. The maximum estimated error is 0.02 pH units. *pKa* values for the mutated proteins different than the ones observed for the wt for shifts larger than the estimated errors are shown in bold. ^***(*)***^ Data obtained from propKa 3 software.^110 (a)^ The predicted values are unusually higher than expected.

|  | *residue* | *propKa*<sup>(*)</sup> | *wt* | *Alpha* | *Beta* | *Gamma* | *Delta* | *Omicron* | *BA.2* | *BA.3* | *BA.4/5* | *BA2.1.21* |
| --- | --- | --- | --- | --- | --- | --- | --- | --- | --- | --- | --- | --- |
| ARG | 346 | 12.43 | 12.34 | 12.35 | 12.35 | 12.35 | 12.31 | 12.36 | 12.35 | 12.35 | 12.33 | 12.35 |
| ARG | 355 | 12.45 | 13.15 | 13.16 | 13.16 | 13.16 | 13.14 | 13.17 | 13.18 | 13.18 | 13.16 | 13.18 |
| ARG | 357 | 12.86 | 12.72 | 12.72 | 12.72 | 12.72 | 12.72 | 12.75 | 12.75 | 12.75 | 12.75 | 12.75 |
| ARG | 403 | 13.33 | 13.90 | 14.00 | 14.00 | 14.00 | 13.88 | **13.70** | **13.31** | **13.29** | **13.29** | **13.31** |
| ARG | 408 | 12.43 | 12.88 | 12.89 | 12.90 | 12.90 | 12.87 | 12.85 | - | **12.64** | - | - |
| ARG | 452 | - | - | - | - | - | **13.48** | - | - | - | **13.46** | - |
| ARG | 454 | 11.30 | 13.62 | 13.63 | 13.62 | 13.62 | 13.54 | 13.55 | 13.54 | 13.54 | 13.53 | 13.54 |
| ARG | 457 | 12.87 | 13.22 | 13.22 | 13.21 | 13.21 | 13.20 | 13.19 | 13.19 | 13.19 | 13.19 | 13.19 |
| ARG | 466 | 11.96 | 13.04 | 13.04 | 13.04 | 13.04 | 13.01 | 13.03 | 13.03 | 13.03 | 13.01 | 13.03 |
| ARG | 493 | - | - | - | - | - | - | **13.13** | **13.11** | **13.11** | - | **13.11** |
| ARG | 498 | - | - | - | - | - | - | **12.81** | **12.78** | **12.78** | **12.78** | **12.78** |
| ARG | 509 | 13.62 | 12.92 | 12.94 | 12.94 | 12.94 | 12.88 | 12.96 | 12.94 | 12.94 | 12.92 | 12.94 |
| ASP | 339 | - | - | - | - | - | - | **3.90** | **3.92** | **3.90** | **3.90** | **3.92** |
| ASP | 364 | 3.56 | 3.95 | 3.96 | 3.95 | 3.95 | 3.95 | 3.97 | 3.98 | 3.97 | 3.97 | 3.98 |
| ASP | 389 | 3.68 | 3.62 | 3.62 | 3.62 | 3.62 | 3.62 | 3.62 | 3.62 | 3.62 | 3.62 | 3.62 |
| ASP | 398 | 6.35<sup>(a)</sup> | 3.16 | 3.16 | 3.16 | 3.16 | 3.13 | 3.14 | 3.16 | 3.14 | 3.15 | 3.16 |
| ASP | 405 | 4.10 | 3.39 | 3.39 | 3.44 | 3.44 | 3.37 | **3.14** | - | - | - | - |
| ASP | 420 | 4.34 | 3.20 | 3.19 | **3.42** | **3.42** | 3.17 | 3.36 | **3.40** | 3.35 | **3.42** | **3.40** |
| ASP | 427 | 3.94 | 3.85 | 3.85 | 3.86 | 3.86 | 3.85 | 3.86 | 3.89 | 3.85 | 3.88 | 3.89 |
| ASP | 428 | 3.48 | 3.97 | 3.98 | 3.98 | 3.98 | 3.98 | 3.98 | 3.99 | 3.98 | 3.99 | 3.99 |
| ASP | 442 | 3.15 | 3.44 | 3.44 | 3.45 | 3.45 | 3.39 | **3.14** | **3.14** | **3.13** | **3.11** | **3.14** |
| ASP | 467 | 2.94 | 3.09 | 3.10 | 3.13 | 3.13 | 3.01 | 3.05 | 3.07 | 3.05 | 3.05 | 3.07 |
| GLU | 340 | 4.53 | 4.05 | 4.05 | 4.05 | 4.05 | 4.04 | **4.29** | **4.29** | **4.29** | **4.29** | **4.29** |
| GLU | 406 | 5.77 | 3.68 | 3.68 | **3.92** | **3.92** | 3.62 | 3.76 | 3.71 | **3.46** | 3.71 | 3.71 |
| GLU | 465 | 3.89 | 3.58 | 3.58 | 3.60 | 3.60 | 3.55 | 3.58 | 3.59 | 3.57 | 3.58 | 3.59 |
| GLU | 471 | 4.61 | 4.20 | 4.20 | 4.18 | 4.18 | 4.10 | 4.10 | 4.10 | 4.10 | 4.11 | 4.10 |
| GLU | 484 | 4.77 | 4.38 | 4.38 | - | - | - | - | - | - | - | - |
| GLU | 516 | 4.29 | 4.05 | 4.06 | 4.06 | 4.06 | 4.06 | 4.06 | 4.06 | 4.06 | 4.06 | 4.06 |
| HIS | 505 | - | - | - | - | - | - | **5.63** | **5.80** | **5.80** | **5.61** | **5.80** |
| HIS | 519 | 6.41 | 6.30 | **5.50** | **5.40** | **5.40** | 6.40 | **5.90** | **6.00** | **5.99** | 6.30 | **6.00** |
| LYS | 356 | 10.38 | 10.63 | 10.63 | 10.63 | 10.63 | 10.62 | 10.72 | 10.72 | 10.71 | 10.71 | 10.72 |
| LYS | 378 | 10.65 | 11.07 | 11.07 | 11.08 | 11.08 | 11.06 | 11.07 | 11.11 | 11.04 | 11.12 | 11.11 |
| LYS | 386 | 10.40 | 11.35 | 11.35 | 11.35 | 11.35 | 11.35 | 11.35 | 11.36 | 11.35 | 11.36 | 11.36 |
| LYS | 417 | 10.12 | 11.39 | 11.40 | - | - | 11.34 | - | - | - | - | - |
| LYS | 424 | 10.32 | 11.36 | 11.36 | 11.39 | 11.39 | 11.34 | 11.37 | 11.40 | 11.35 | 11.40 | 11.40 |
| LYS | 440 | - | - | - | - | - | - | **10.58** | **10.57** | **10.56** | **10.57** | **10.57** |
| LYS | 444 | 10.34 | 10.76 | 10.81 | 10.81 | 10.81 | 10.74 | **10.58** | **10.57** | **10.56** | **10.56** | **10.57** |
| LYS | 458 | 10.51 | 10.59 | 10.58 | 10.59 | 10.59 | 10.54 | 10.54 | 10.55 | 10.54 | 10.56 | 10.55 |
| LYS | 462 | 10.56 | 10.95 | 10.95 | 10.96 | 10.96 | 10.94 | 10.96 | 10.96 | 10.95 | 10.96 | 10.96 |
| LYS | 478 | - | - | - | - | - | **10.67** | **10.67** | **10.67** | **10.67** | **10.67** | **10.67** |
| LYS | 484 | - | - | - | **10.60** | **10.60** | - | - | - | - | - | - |
| TYR | 351 | 10.54 | 9.60 | 9.59 | 9.61 | 9.61 | **9.29** | 9.55 | 9.55 | 9.54 | **9.33** | 9.55 |
| TYR | 365 | 12.13 | 10.10 | 10.10 | 10.10 | 10.10 | 10.09 | 10.14 | 10.14 | 10.13 | 10.14 | 10.14 |
| TYR | 369 | 11.41 | 9.72 | 9.72 | 9.72 | 9.72 | 9.71 | 9.73 | 9.73 | 9.72 | 9.73 | 9.73 |
| TYR | 380 | 10.43 | 9.64 | 9.64 | 9.64 | 9.64 | 9.63 | 9.64 | 9.67 | 9.63 | 9.67 | 9.67 |
| TYR | 396 | 10.24 | 9.42 | 9.42 | 9.42 | 9.42 | 9.41 | 9.44 | 9.44 | 9.44 | 9.44 | 9.44 |
| TYR | 421 | 10.83 | 9.48 | 9.48 | **9.72** | **9.72** | 9.44 | **9.66** | **9.67** | **9.64** | **9.70** | **9.67** |
| TYR | 423 | 16.31<sup>(a)</sup> | 9.64 | 9.64 | 9.69 | 9.69 | 9.62 | 9.68 | 9.71 | 9.66 | 9.70 | 9.71 |
| TYR | 449 | 10.20 | 9.71 | 9.73 | 9.73 | 9.73 | 9.61 | **9.47** | **9.47** | **9.45** | **9.44** | **9.47** |
| TYR | 451 | 12.58<sup>(a)</sup> | 9.86 | 9.88 | 9.89 | 9.89 | **9.59** | 9.77 | 9.77 | 9.76 | **9.53** | 9.77 |
| TYR | 453 | 10.85 | 9.68 | 9.69 | 9.95 | 9.95 | **9.33** | **9.33** | **9.31** | **9.27** | 9.56 | **9.31** |
| TYR | 473 | 10.30 | 9.63 | 9.63 | 9.60 | 9.60 | 9.46 | 9.45 | 9.45 | 9.45 | 9.48 | 9.45 |
| TYR | 489 | 10.04 | 9.87 | 9.87 | 9.73 | 9.73 | 9.72 | 9.68 | 9.67 | 9.68 | 9.74 | 9.67 |
| TYR | 495 | 14.87 | 10.03 | 10.07 | 10.14 | 10.14 | **9.74** | 9.84 | **9.78** | **9.75** | **9.64** | **9.78** |
| TYR | 501 | - | - | **9.81** | **9.82** | **9.82** | - | **9.25** | **9.20** | **9.18** | **9.20** | **9.20** |
| TYR | 505 | 10.15 | 9.70 | 9.86 | 9.89 | 9.89 | 9.69 | - | - | - | - | - |
| TYR | 508 | 10.22 | 9.65 | 9.67 | 9.70 | 9.70 | 9.64 | 9.58 | 9.51 | 9.45 | 9.51 | 9.51 |

Since most of the previous simulation studies adopted propKa^110^ to define the protonation states of the RBD to set-up the charges for their simulations, we included the *pKa* values obtained from propKa for the RBD_(wt)_. The root-mean-square deviations (RMSD) between the results predicted by the FPTS and the ones from “propKa 3.0”^110^ are equal to 1.57 pH units, slightly higher than what was found for a benchmark with a larger set of proteins.^88^ Some *pKa* values from propKa are unusually higher than what would be expected (e.g. the pKa for D398 was 6.35, a shift of 2.35 pH units from its ideal value). Excluding these unrealistic values, the RMSD drops down to 0.95.

### Electrostatic coupling between titratable residues

The presence of a large number of ionizable groups at the RBD-ACE2 interface and the electrostatic features presented above suggest a high number of coupled titratable sites due to the long-range nature of the electrostatic interactions. This was mapped by monitoring the residues affected by the replacement of an ALA. For each single ALA mutation, a full titration study was performed computing the *pKas* of all other ionizable groups for this mutant RBD protein. Comparing the results with the ones obtained for the wt, it was possible to identify all amino acids whose *pKas* were shifted (Δ*pKa*_*i*_), i.e. the ones electrostatically coupled with the replaced residue. Mathematically, for each titratable amino acid *i* in the RBD protein with the ALA mutation, it was calculated Δ*pKa*_*i*_=*pKa*_*i*_(mutant)-*pKa*_*i*_(wt), where *pKa*_*i*_(wt) is the *pKa* value of the amino acid *i* for the RBD_(wt)_ (the values are listed in Table 2) and *pKa*_*i*_(mutant) is its value for the mutant RBD. For example, when GLU-340 is mutated by ALA (E340A), six other titrable groups are affected (CYS-336, LYS-356, ARG-357, CYS-361, ARG-509, and HIS-519). The number of electrostatically coupled amino acids with a specific one (*N*_*coupled*_) is given in Figure 3 for all titratable groups of the RBD_(wt)_. On average, a titratable residue in the RBD_(wt)_ is electrostatically coupled with 6.8 others (<*N*_*coupled*_>=6.8±2.7). The cases with higher *N*_*coupled*_ values are distributed over the RBD surface as can be illustrated in Figure 1. Three amino acids (ASP-420, ARG-454, and ARG-457) have *N*_*coupled*_ equals 12, and one (ASP-467) has *N*_*coupled*_ equals 13. The latter, i.e. the residue ASP-467, is electrostatically connected to TYR-351, TYR-421, TYR-423, LYS-424, ARG-454, ARG-457, LYS-458, LYS-462, GLU-465, ARG-466, GLU-471, TYR-473, and HIS-519. It is worth noting that not only neighbor residues are affected but also amino acids that are spatially distant (e.g. the distance ASP-467-HIS-519 is ∼32Å). This shows that mutations at the RBD-ACE2 interface may impact other parts of the protein and vice versa. There are also exceptions. For example, although HIS-519 is often affected by the neutralization of other titratable groups, the replacement of HIS-519 by ALA (H519A) only affects GLU-516 [ *N*_*coupled*_(HIS-519)=1] that is located in its neighborhood. It is easier to see how all titratable groups are electrostatically connected in a simplified correlation network diagram as seen in Figure 4.

**Figure 3:**
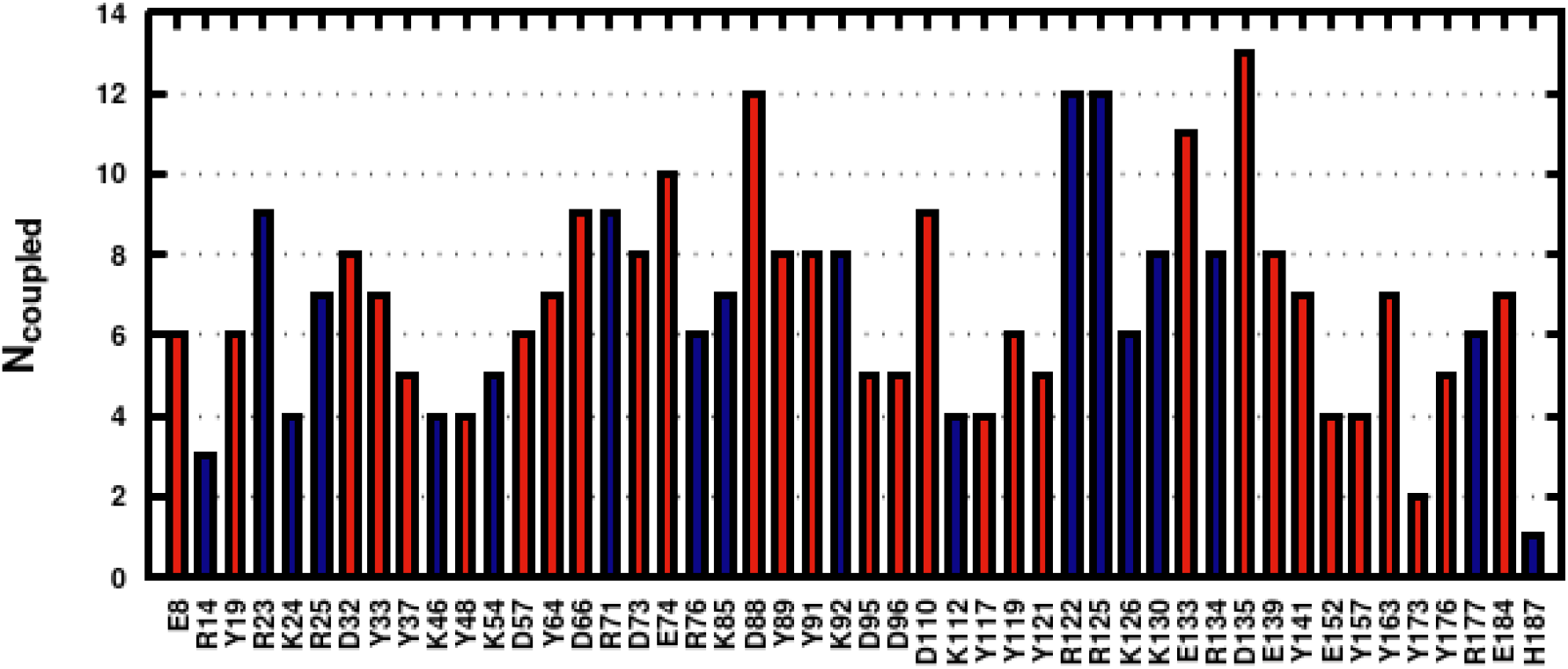
Number of amino acids electrostatically coupled with others per titratable amino acid. Amino acids are given in the one-letter format with their numbers displayed in an inner numbering system. The corresponding numbers in the PDB system are obtained by adding 332 (e.g. E8 is E340/GLU-340). See text for more information.

**Figure 4:**
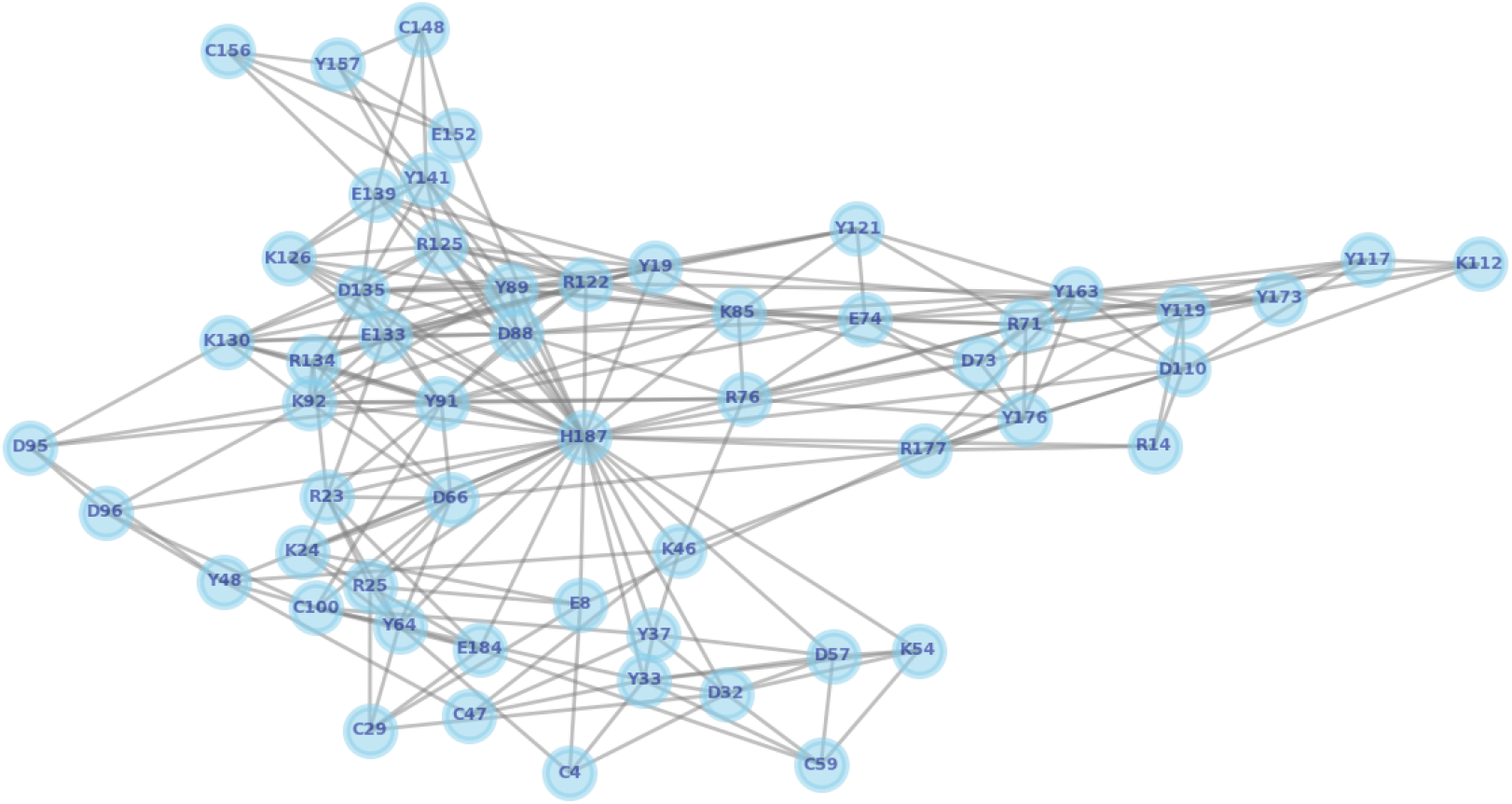
A simplified correlation network diagram in two-dimension for the electrostatic coupling of the titratable groups. The diagram was built-up with all the titratable groups as nodes and the other residues that they affect as connections based on the *pKa* analysis as described in the text. Amino acid labels are given as in Figure 3.

A similar analysis was done for the VOCs and the sub-lineages of Omicron - see Figure 5. Instead of *pKa*’s from a single ALA mutation, the data from Table 2 was used for both *pKa*_*i*_(mutant) and *pKa*_*i*_(wt). It is intriguing to see that the identified clusters follow the phylogenetic tree^111^ indicating again that the viral evolution is intimately connected with the electrostatic properties of its proteins dictating the viral physiopathology. This result generalizes a recent analysis done at pH 7 and using electrostatic surfaces of RBD proteins for different variants showing an equivalent interpretation.^62^ The number of titratable residues affected by each set of mutations of the variants also increased over time (since the Wuhan outbreak) and their transmissibility. The smaller cluster is found for the Beta/Gamma variants (that appeared in the first year of the pandemic) essentially with *N*_*coupled*_ equals to 6 residues. The Delta variant is more transmissible and has an intermediate situation with *N*_*coupled*_ equals to 8 residues. It is also an intermediate variant in terms of its surge time after Beta/Gamma and before Omicron. The larger cluster is seen for the Omicron family (that were detected more recently during the last months) which shares several affected amino acids in common. The mutations found in Omicron affect other 17 amino acids while BA.3 and BA.4/BA.5 have the higher *N*_*coupled*_ value (*N*_*coupled*_=19).

**Figure 5:**
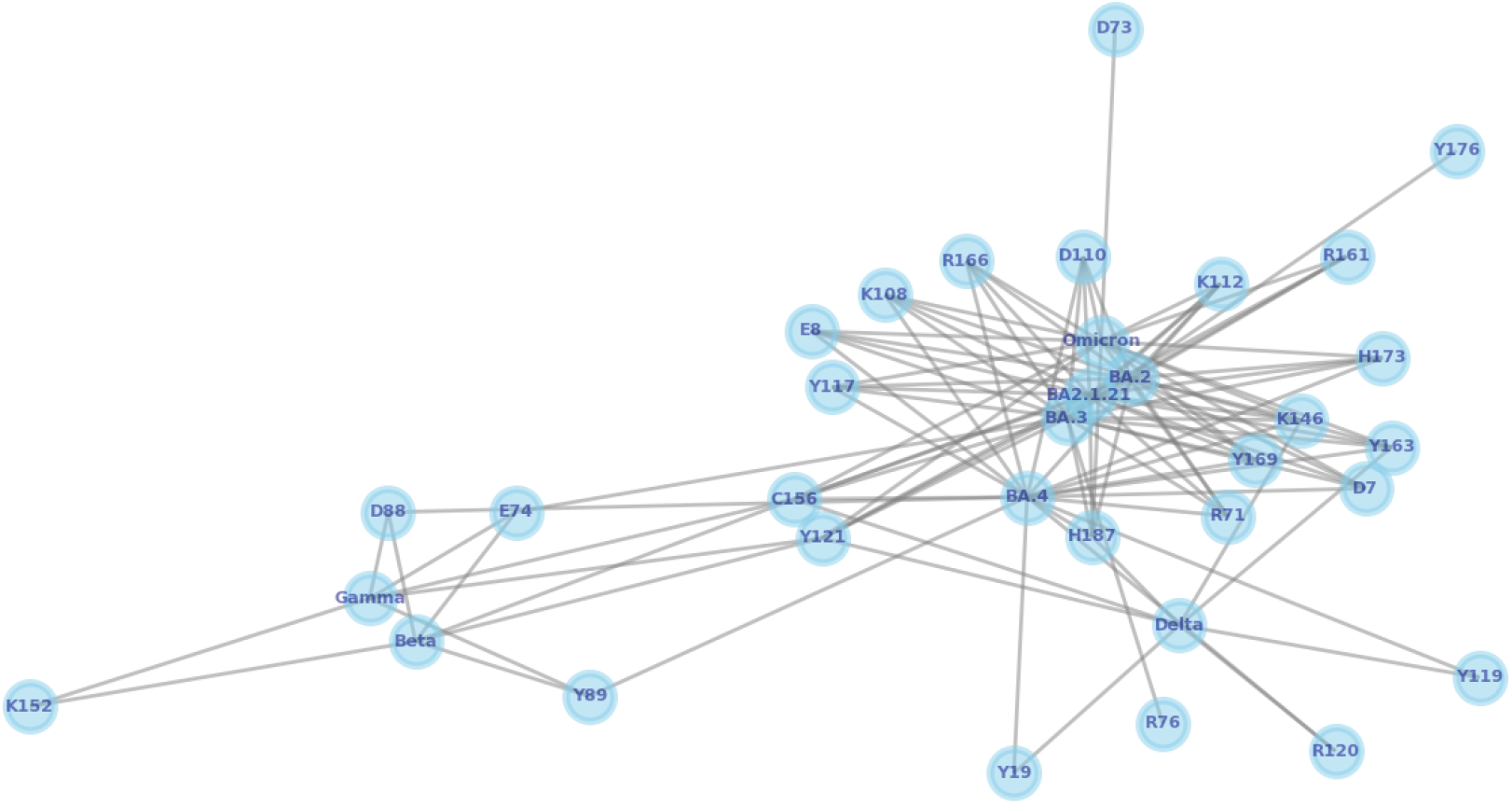
A simplified correlation network diagram in two-dimension for the electrostatic coupling of the titratable groups for the mutations present in different SARS-CoV-2 variants. The diagram was built-up with the analyzed variants as nodes and the affected amino acids as connections based on the *pKa* data given in Table 2. Amino acid labels are given as in Figure 3.

### Free energy of interactions at pH 7

The physical-chemical parameters discussed above already indicate that the RBD-ACE2 attraction is increasing for the new variants. This was confirmed by the estimated free energy of interactions obtained by the FORTE approach. From the computed plots of *w(r)* as a function of the separation distance between the RBD proteins and ACE2 at pH 7 and 150mM of salt, we extracted the well-depth values (*w*_*min*_) for each studied system to facilitate the analysis. Figure 6 summarizes the main comparisons for the wild-type and several variants together with the total net protein charge numbers. The negative values of the computed *w*_*min*_ indicate the complexation for all the systems. It can be seen that the charge and the binding affinities are increasing for the VOCs (Alpha, Beta, Gamma, Delta, and Omicron). Other VOIs and mutations can be arranged within the approximations of the used model in the same groups, from the lower to the higher predicted complexation affinities: **a)** Alpha (wt, B.1.640.1/Fr, B.1.640.2-IHU/FR, and mink), **b)** Beta/Gamma (Epsilon and Mu), **c)** Eta (Kappa, Iota, Mu [without the K417T mutation], a double L452R and T478K mutations), **d)** Delta (no other variant together), **e)** Omicron (BA.2, BA.4/BA.5 and BA.2.12.1), and **f)** BA.3. The group of Eta does not include a VOC but it was found between two groups with VOCs (Beta/Gamma and Delta). The mutation E484K present in Eta, Iota, and Mu (among others) was found to improve its binding affinity for ACE2 (as predicted here too) and to be critical for the virus.^36^ BA.3 is also isolated from the other VOCs and, as seen for Z and *μ*, it has an outstanding behavior. The computed values for the *w*_*min*_ values are consistent with the physical-chemical parameters reported above. Comparisons of the binding affinities for the VOCs with other theoretical studies were reported by us in previous work.^23^ Other results that became available after this comparison also report the same trends.^61,62^ For the recent variants from the Omicron family, no data is yet available at the moment of writing this work.

**Figure 6:**
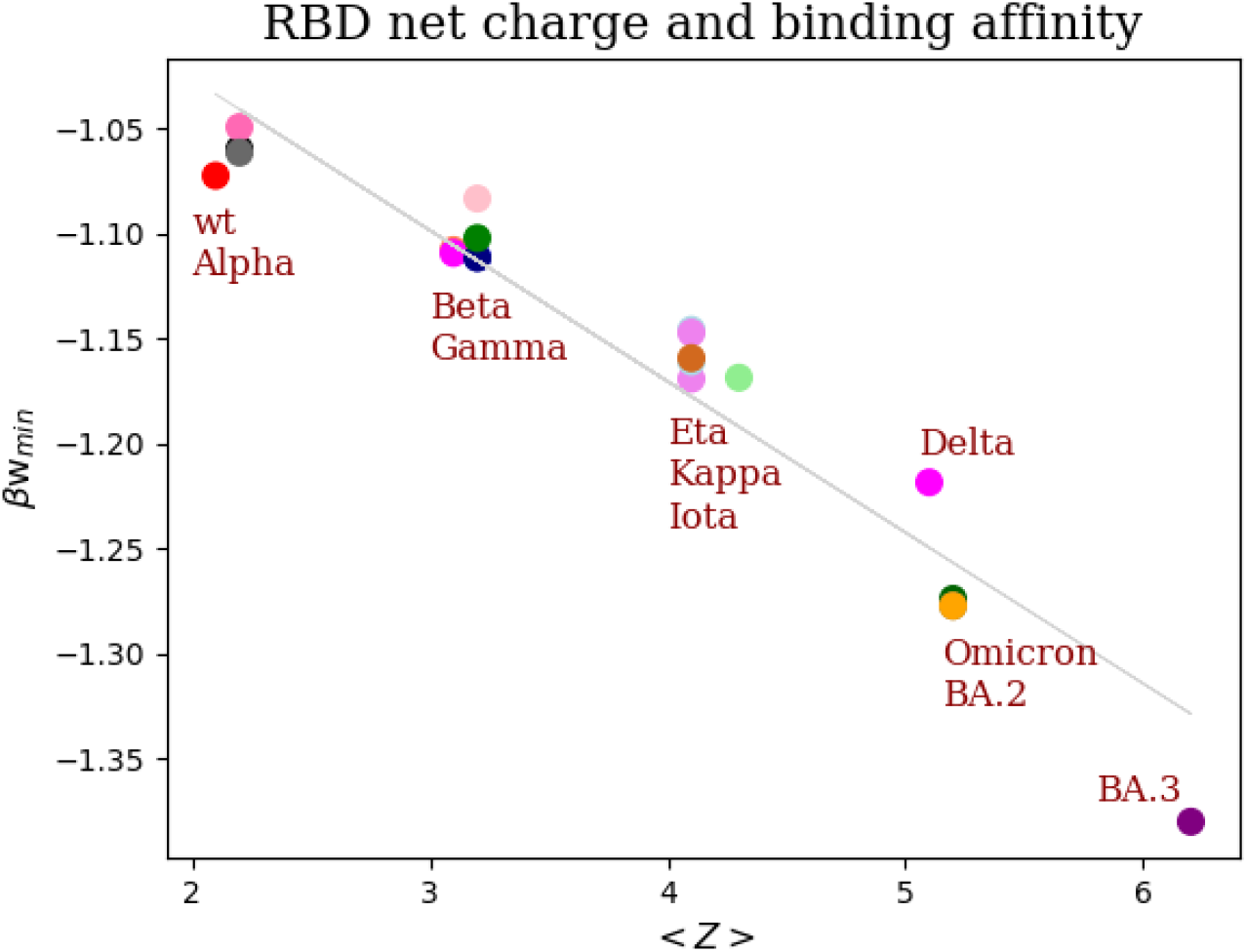
Correlation between the total net charge number of each RBD and the corresponding βw_min_ values for their complexations with ACE2. The data for the total net charge numbers were obtained from titration simulations with a single protein in the absence of the ACE2 by the FPTS at pH 7 and 15mM of salt. All βw_min_ values (given in K_B_T units) were computed from RBD-ACE2 complexation studies using FORTE at pH 7 and 150mM of NaCl. The estimated error is 0.01 K_B_T. The gray line is a linear regression plot obtained using only the five VOCs as the training set (βw_min_= -0.073525*<Z> -0.88973, regression coefficient =-0.96). See text for more details.

Different independent factors can contribute to increasing virus transmission, infectivity, and severity.^23,32,58,112,113^ The RBD-ACE2 binding affinity controls the SARS-CoV-2 entry mechanism that allows the infection to occur.^37^ This makes it an important factor among the others. However, a stronger RBD-ACE2 binding affinity also relies on at least other two interconnected mechanisms for an optimal cell invasion by the virus: **a)** the S protein has to be at the “up” conformational state to permit the RBD to be exposed to the ACE2,^32^ and **b)** an efficient interaction between the S protein with the host protease furin for its cleavage.^113^ Mutations can have a direct or indirect effect on all of them. Some studies have suggested a direct connection between a stronger binding affinity with increased transmissibility.^23,105,107^ In fact, transmission has increased over time and the surge of the new variants. For example, Omicron is more transmissible than Delta^114^ and it has a stronger RBD-ACE2 binding affinity in our computer data. The analysis of the transmission in Danish Households revealed that BA.2 is more transmissible than Omicron.^115^ However, in our calculations, the difference in their binding affinities is within the statistical significance: *w*_*min*_(Omicron)=-1.27(1) and *w*_*min*_(BA.2)=-1.28(1). The same situation is verified for the Alpha variant. The reverse genetics study carried out by Liu and colleagues showed an increased transmission for Alpha in comparison to the wt.^116^ The computed statistical deviation does not allow us to make such a statement: *w*_*min*_(wt)=-1.06(1) and *w*_*min*_(Alpha)=-1.07(1). These discrepancies could be due to several reasons: **i)** intrinsic approximations of the simulated model, **ii)** the need to include the other factors mentioned above in the simulations, and **iii)** the apparent lack of consensus in the experimental data.^117^ The latter issue has been recently discussed in the literature.^23,63,117^

On one hand, the model can capture larger discrepancies between different variants as for the comparison between Delta and Omicron. On the other hand, differences for a single mutation can be within the statistical significance even when the reported experimental assays show a larger enhancement for the binding affinity. Bayarri-Olmos et al. reported a biolayer interferometry analysis demonstrating a 4-fold higher affinity for the mink variant than the wt strain (*K*_*D*_ = 15.5 nM for RBD_(wt)_-ACE2 and *K*_*D*_ = 3.85 nM for RBD_(mink)_-ACE2, where *K*_*D*_ is the equilibrium dissociation constant).^107^ In our calculations, the predicted affinity is identical: *w*_*min*_(wt)=-1.06(1) and *w*_*min*_(mink)=-1.06(1). Conversely, it was reported a quite dispersed experimental data for the RBD_(wt)_-ACE2 binding affinity: *K*_*D*_ = ∼1.2–14.7 nM (see discussion in refs. ^32,117^) which makes it difficult for such analysis at the level of binding affinities. It becomes more challenging when we try to establish the link between affinity and transmissivity. The computed data for the RBD_(BA.3)_-ACE2 would suggest that BA.3 should be more transmissible than the other sublineages from Omicron’s family. However, this variant surprisingly did not show such expected higher transmission and is not often detected in the genomic surveillance centers (the Network for Genomic Surveillance in South Africa reported in May/2022 that BA.3 is detected at low levels [0.14% - see https://www.nicd.ac.za/]). The apparent puzzle is solved when structural data for the whole S trimer reveals that the S protein adopts closed- or semi-closed forms for the BA.3 preventing the RBD to complex with ACE2.^105^ Taken together, it is safe to say that the RBD-ACE2 binding affinity is a good predictor to estimate the *tendencies* of a high transmissible variant and increased infectivity. As long as mutations do not have a substantial impact on the up state of the S protein or in its interactions with furin or with any other molecule that can add to the entering process (e.g. sialic acid^118^), the predictions of the transmissivity using the binding affinities will be reliable. A possible limitation would be that external factors such as the mobility of the contaminated people that can contribute to the transmission of the virus are not incorporated into the present framework.

Based on this analysis, we can speculate that such a high binding affinity computed for the Omicron family is strong enough to allow their RBDs to complex faster and tightly with ACE2 before Abs (produced by the humoral immune system or administered as therapeutics) can detect, identify and/or block it. This could be a possible rational explanation for the difficulties to prevent reinfections and new breakthroughs of vaccinated people, and the short-lasting period of protection for Omicron’s contaminated patients.^97^ Thinking on the molecular aspects, it might be more challenging for the B cells to form an immune memory of these RBD proteins if they do not have enough time to interact with the RBD due to their strong RBD-ACE2 binding affinities. Some additional support for this possible explanation is the fact that the vaccine efficacy against infection is apparently determined by the Ab binding in the S protein to block its interaction with ACE2.^119^

### A simple predictor for RBD-ACE2 binding affinity

The linear regression plot (lrp) calculated for the <*Z*>s of the RBD proteins and the βw_min_s using the corresponding computed values for Alpha, Beta, Gamma, Delta, and Omicron variants (shown in Figure 6) suggests that it would be possible to predict the RBD-ACE2 binding affinity for a new RBD solely using its net charge value. We tested this simple “charge rule” predicting *βw*_*min*_ for the other variants (not included in the training dataset) at pH 7 and 150 mM of salt using their values of <*Z*> from titration studies at the same physical-chemical condition. The comparison between the measured (given by the points in Figure 6) and predicted values from the lrp resulted in an RMSD of 0.03 K_B_T units. For simpler mutations as single replacements of a titratable group by an alanine (see the data in the next subsection), this linear model performs better with an RMSD of 0.004 K_B_T units (all 48 ALA mutations from Figure 7 were included in this test dataset) which is smaller than the estimated errors for *βw*_*min*_.

**Figure 7:**
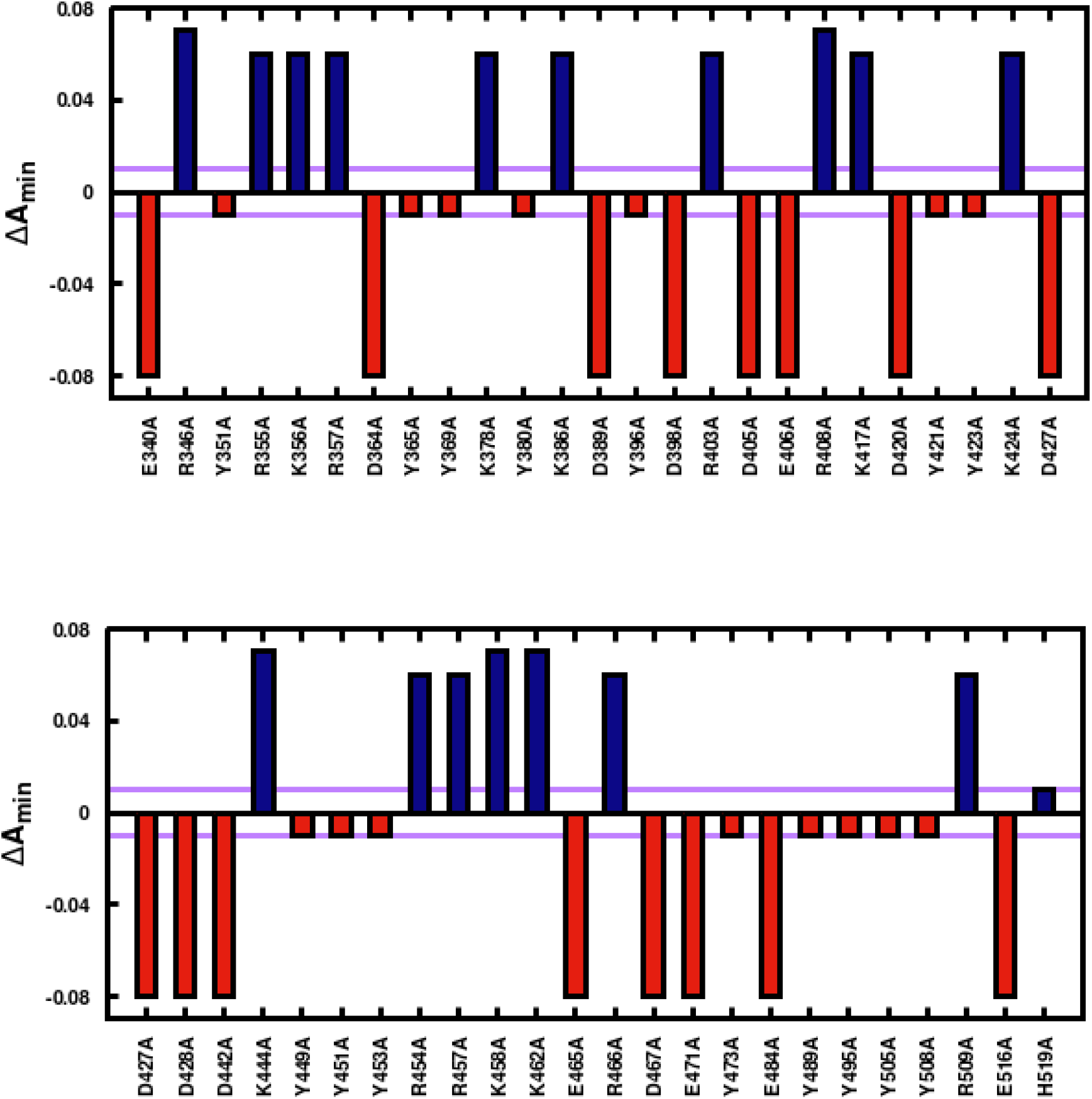
Individual residue electrostatic contributions to the complexation SARS-CoV-2 RBD-ACE2. Acid and basic amino acids are represented by red and blue, respectively. Data from the theoretical ALA scanning procedure applied to the RBD_(wt)_. ΔA_min_ (in K_B_T units) is defined as the difference between the minimum value measured in βw(r) for the complexation RBD_(mutated)_-ACE2 and the corresponding quantity for the RBD_(wt)_-ACE2. The purple line indicates the estimated statistical error in the computed *βw*_*min*_ values. See the text for more details.

The slope of the lrp (−0.073525) is numerically in the same order as the Coulombic contributions estimated from a simple Derjaguin–Landau–Verwey–Overbeek (DVLO)-based analytical approach

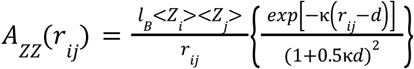

where *l*_*B*_ is the Bjerrum length and *d* an adjustable parameter.^109^ For example, for the Epsilon variant (<*Z*_(Eta)_>=3.1, <*Z*_(ACE2)_>=-23.1, *l*_*B*_=7.1Å, *r*_*ij*_=49Å, κ=7.9Å, and *d*=20Å), *A*_*ZZ*_*(r*_*ij*_) equals to -0.05 K_B_T. The average value of *A*_*ZZ*_*(r*_*ij*_) for all variants used in the validation procedure is -0.06 K_B_T. The small difference between this value and the slope of lrp is from the other electrostatic terms.^109^ The dominant contribution is from the charge-charge term as suggested before.^9,10,23,32^ ACE2 polymorphisms follow the same rule.^23^ The intercept (−0.88973) is an estimate of the non-electrostatic attractive contributions. This simple “charge rule” will work for future RBD proteins provide their dipole numbers are not so different from the ones seen in Figure 2d. It is a practical means to quickly *estimate* the RBD-ACE2 affinity and the virus transmissivity and infectivity tendencies. More precisely analytical calculations can also be performed.^109,120^

### Key amino acids for the complexation and their contributions

RBD-ACE2 complexation studies were performed with man-created RBD proteins obtained with the replacement of a titratable residue by an ALA. All calculations for this purpose were carried out at pH 7 and 150mM of salt. Using this theoretical ALA scanning procedure, we could identify the contributions of each particular amino acid to the estimated free energy of interactions. The template for all mutations was the RBD_(wt)_, and all the 48 titratable groups were evaluated in 288 MC simulation runs [48 mutants*3 replicates*(1 equilibration + 1 production)]. Due to the computational costs of this large set of runs, only a single replacement per system was explored and all tests were performed solely for the RBD_(wt)_. Figure 7 displays the computed *ΔA*_*min*_ defined as the difference between *βw*_*min*_(mutant) and *βw*_*min*_(wt) [*ΔA*_*min*_*=βw*_*min*_(wt)-*βw*_*min*_(mutant)] under the same physical-chemical conditions. Mutations that favor the complexation are shown in red (*ΔA*_*min*_*<0*) while the ones that diminish the RBD-ACE2 binding affinity are given in blue (*ΔA*_*min*_*>0*). As expected, the replacement of a negatively charged amino acid by ALA (e.g. E340A) increases < *Z*(RBD)> from +2.2 (measured for the wt) to +3.1 which favors the Coulombic attraction with the negatively charged ACE2 [<*Z*_(ACE2)_>=-23.1(1)]. Conversely, mutating a positively charged residue by ALA works in the opposite direction: R346A has a smaller net charge [<*Z*_(R346A)_>=+1.2(1)] in comparison to the RBD_(wt)_ reducing the charge-charge attractive term. In summary, mutations that result in a more positive RBD favor its binding to ACE2.

### pH effects on the RBD-ACE2 complexation

Computed free energy of interactions for the RBD-ACE2 complexation for all studied variants at 150mM of salt and several pH values are given in Figure 8. For the sake of convenience, Figure 8a displays the compiled *βw*_*min*_ values for all systems using a heatmap-style plot. The corresponding estimated *βw*_*min*_ values are given inside the colored boxes. The darkest green is the system and the pH condition with the highest predicted RBD-ACE2 binding affinity. Conversely, the darkest red is given to the system/pH condition with the lowest predicted binding affinity. The same trends discussed in the previous subsection specifically for pH 7 are repeated at all pH regimes. This confirms that predictions obtained at pH 7 can be qualitatively used to understand the behavior of the system at the other pH regimes. Complexation can happen in a larger pH window which suggests that the SARS-CoV-2 entering mechanism is not pH-dependent. However, depending on the medium and clinical condition of the patient, the complexation can be tighter. All systems have the maximum attraction around pH 7.5. This is straightforwardly seen in Figure 8b which gives a complementary view by showing the *βw*_*min*_ values as a function of pH for some key systems (wt, Delta, and Omicron). It illustrates how the mutations found in the variants favor the RBD-ACE2 complexation following the temporal order these VOCs appeared and circulated. The charge-charge contribution captured by the dashed line indicates the dominance of this electrostatic term in particular around pH 7.

**Figure 8:**
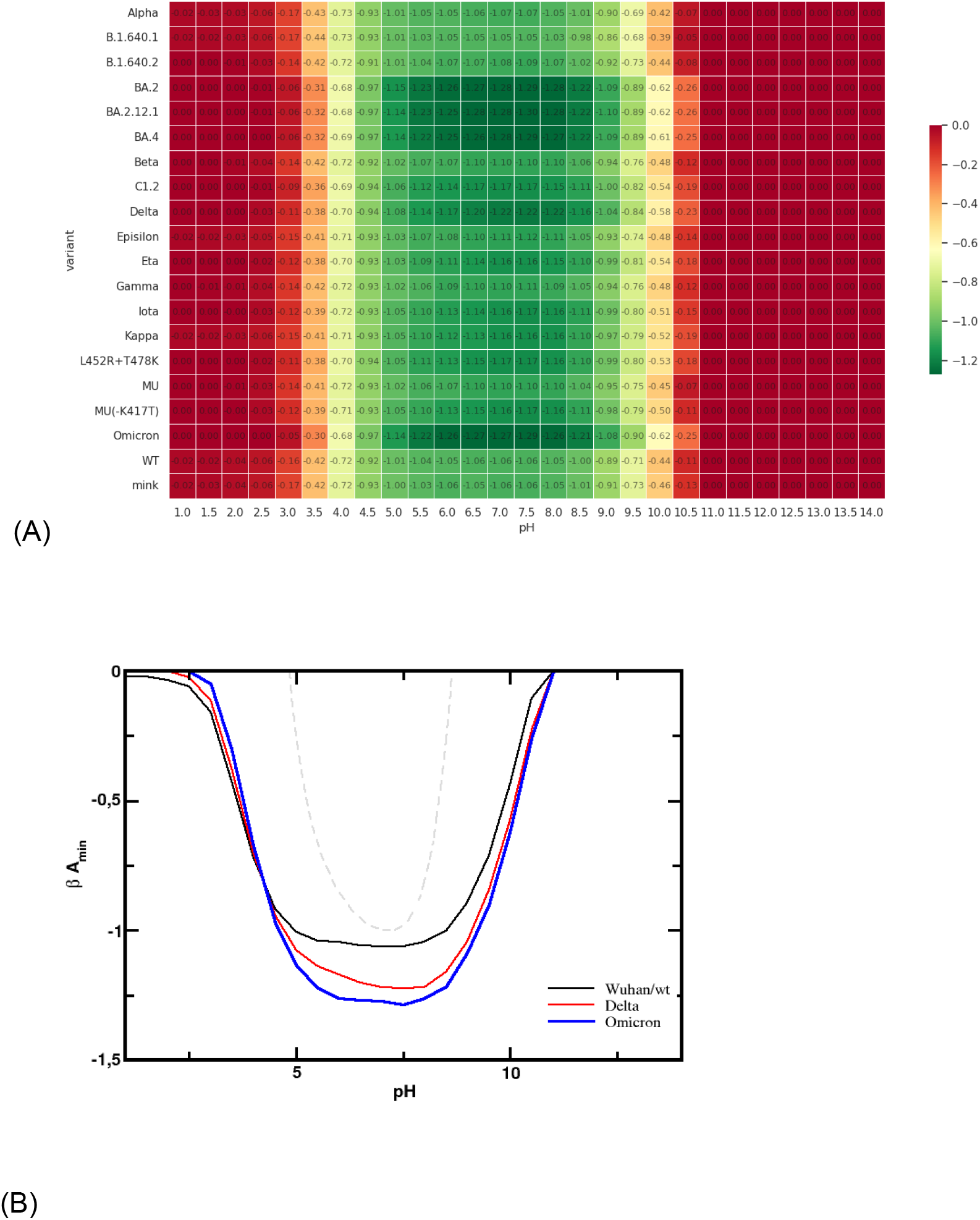
pH effect on the simulated free energy of interactions for the RBD-ACE2 complexation. All simulations were performed at 150 mM of salt. All values of *βw*_*min*_ are given in K_B_T units. The maximum estimate error is 0.01. **(a)** Heatmap with the minima free energy of interactions values measured for the SARS-CoV-2 RBD-ACE2 complexation at different pH conditions for the main variants. The computed *βw*_*min*_ values are given inside each box. **(b)** *βw*_*min*_ values as a function of the pH solution for some selected cases (wt, Delta, and Omicron variants). The dashed line is the normalized <*Z*_(RBDwt)_>*<*Z*_(ACE2)_> at different pH conditions. The minimum value of the product was used to divide all the computed points.

## 4. Conclusions

Electrostatic interactions can be used in a variety of ways by viruses. We have shown that SARS-CoV-2 RBD is evolutionarily becoming more and more positively charged which favors its computed binding affinity to ACE2. The predicted binding can be observed in a broad pH window although it is stronger at pH 7.5. The COVID19 virus is also gradually increasing the dipole moment of the RBD of the new variants to improve further its binding affinity to enter the human cells. The RBD-ACE2 association is mainly controlled by Coulombic charge-charge interactions for the known variants. The charge regulation mechanism plays a minor role in this complexation process. This result provides a more solid physical basis for simulations using only the constant-protonation state approach. Based on the present analysis, a predictor for virulence and transmitivity was devised using titration properties. This tool can help to identify the human risks of the variants that will appear in the near future. In addition, we have shown the high electrostatic coupling between the titratable groups. This is an important result to illustrate that ionizable groups outside the biological RBD-ACE2 interface are also important for the complexation mechanism. The long-range nature of the electrostatic interactions seen in the present analysis reinforces the limitations of the traditional “lock-and-key” model for molecular complexation. Different electrostatic features such as the pKas correlate with phylogenic analysis connecting physical chemistry with the virus evolution. Our results contribute to the general physicochemical understanding of the biomolecular electrostatic phenomena and shed light on the virus evolution and its molecular mechanisms pH-dependent.

## ACKNOWLEDGMENTS

This work has been supported in part by the “Fundação de Amparo à Pesquisa do Estado de São Paulo” [Fapesp 2020/07158-2 (F.L.B.d.S.)] and the Conselho Nacional de Desenvolvimento Científico e Tecnológico (CNPq) [CNPq 305393/2020-0 (FLBdS)]. F.L.B.d.S. is also deeply thankful for resources provided by the Swedish National Infrastructure for Computing (SNIC) at NSC and PDC. A. Laaksonen acknowledges the Swedish Research Council for financial support, and partial support from a grant from the Ministry of Research and Innovation of Romania (CNCS - UEFISCDI, project number PN-III-P4-ID-PCCF-2016-0050, within PNCDI III). This research was also done using services provided by the OSG Consortium, which is supported by the National Science Foundation awards #2030508 and #1836650.

